# Phosphate transporter PstSCAB of *Campylobacter jejuni* is essential for lactate-dependent growth and colonization in chickens

**DOI:** 10.1101/843771

**Authors:** Ritam Sinha, Rhiannon M. LeVeque, Marvin Q. Bowlin, Michael J. Gray, Victor J. DiRita

## Abstract

*Campylobacter jejuni* causes acute gastroenteritis world-wide and is transmitted primarily through poultry, in which it is often a commensal member of the intestinal microbiota. Previous RNASeq experiments showed that transcripts from an operon encoding a high affinity phosphate transporter (PstSCAB) of *C. jejuni* were among the most abundant when grown in chickens. Elevated levels of the *pstSCAB* mRNA were also identified in an RNASeq experiment from human infection studies. In this study, we explore the role of PstSCAB in the biology and colonization potential of *C. jejuni.* Our experimental results demonstrate that cells lacking PstSCAB survive poorly in stationary phase, nutrient-limiting media, and under osmotic conditions reflective of those in the chicken. Polyphosphate levels in the mutant cells were elevated at stationary phase, consistent with alterations in expression of polyphosphate metabolism genes. *C. jejuni* were highly attenuated in colonization of newly hatched chicks, recovered at levels several orders of magnitude below wild type. Mutant and wild type grew similarly in complex media but the *pstSCAB* mutant exhibited a significant growth defect in minimal media supplemented with L-lactate, postulated as a carbon source *in vivo*. Poor growth in lactate correlated with diminished expression of acetogenesis pathway genes previously demonstrated as important for colonizing chickens. The phosphate transport system is thus essential for diverse aspects of *C. jejuni* physiology and *in vivo* fitness and survival.

**Importance:** *C. jejuni* causes millions of gastrointestinal infections annually worldwide. Poultry and poultry products are major sources of *C. jejuni* infection to human as the microbe is a commensal colonizer of the chicken gastrointestinal tract. Due to the emergence of multi-drug resistance in *C. jejuni*, there is need to identify alternative ways to control this pathogen. Genes encoding the high-affinity phosphate transporter PstSCAB were highly expressed during colonization of *C. jejuni* in chicken and human. In this study, we address the role this high-affinity phosphate transporter PstSCAB of *C. jejuni* on chicken colonization and for its general physiology. PstSCAB is required for colonization in chicken, metabolism and survival under different stress responses and during growth on lactate, a potential substrate for growth of *C. jejuni* in chickens. Our study highlights that PstSCAB may be an effective target to develop mechanisms to control the bacterial burden in both chicken and human.

## Introduction

*Campylobacter jejuni* is a microaerophilic, Gram-negative bacterium that causes acute gastroenteritis worldwide (1). Common campylobacteriosis symptoms are self-limiting diarrhea (which may be watery to bloody), with nausea, fever and abdominal cramping in human. As *C. jejuni* is a common member of the chicken gastrointestinal microbiota, poultry meat is a major source of human campylobacteriosis, either through direct consumption of poultry or through cross-contamination of other food (2, 3). As consumption of poultry meat worldwide over the last 20 years has approximately doubled, (4) the threat of campylobacteriosis remains high.

To explore mechanisms by which *Campylobacter* colonizes the chicken gastrointestinal tract, we used RNASeq to determine the transcriptome of *C. jejuni* strain 81-176 harvested from ceca of seven day-old chickens after experimental oro-gastric inoculation on day-of-hatch (5). Among the more highly abundant transcripts in this population were those encoding a predicted phosphate transporter (*pstSCAB*), suggesting that phosphate transport is important for the *in vivo* fitness of *C. jejuni*, and perhaps that inorganic phosphate plays a key role of chicken colonization (5).

Phosphorus, or inorganic phosphate (Pi), is one of the most important macronutrients for microbial growth. It is required for biosynthesis of cellular components such as nucleic acid, phospholipids and proteins, and is critical for maintaining a cellular energy source in ATP. It is also involved in regulating carbon and nitrogen metabolism (6). Uptake and storage of inorganic phosphate, a growth-limiting nutrient, is heavily regulated in bacteria. Generally, in Gram-negative bacteria, a two-component sensor-kinase/response regulator system controls phosphate transport in response to levels of available inorganic phosphate. In *C. jejuni*, a two component regulatory system, PhosSR, controls the high affinity phosphate transporter operon (*pstSCAB*), as well as other genes, in phosphate limiting condition (7). PstSCAB comprises an ATP binding cassette (ABC) transporter system in the inner membrane of many Gram-negative bacteria, with PstS the periplasmic phosphate binding protein, PstAC comprising transmembrane protein permeases and PstB an ATP binding and hydrolyzing subunit. PstS bound to Pi is stimulated to associate with PstA/C transmembrane proteins. This triggers ATP hydrolysis in the PstB dimer which induces the conformational changes that facilitate release of Pi into the cytosol (8). In many bacteria a PstSCAB-associated protein, PhoU, acts to reduce expression of the Pho regulon under conditions of phosphate sufficiency (6), but a PhoU homologue has not been reported in *C. jejuni*.

Phosphate transporters contribute to virulence in other microbes, although the molecular mechanisms by which that occurs varies across species and in any event are incompletely understood (9). For example, inoculation into chickens of an avian pathogenic *E.coli* mutant strain lacking *pst* led to fewer extra-intestinal lesions than did inoculation of wild type, and the mutant also showed higher sensitivity to rabbit serum (though not to chicken serum), polymyxin B, and acid shock (10). A *pstS* mutant of virulent EHEC and EPEC showed significant adherence and colonization defects in intestinal epithelial cells as well as in animal models, resulting from altering expression of adhesin proteins (9, 11). *Mycobacterium tuberculosis* contains two different phosphate transporters which regulate replication in macrophages and provide protection from antimicrobial stress response (12, 13).

Given the high abundance of transporter mRNA during *in vivo* growth of *C. jejuni*, it seems likely that the transporter is a determinant of colonization. We analyzed a mutant lacking *pstSCAB* for several growth phenotypes as well as chicken colonization. We demonstrate a severe colonization defect in mutants lacking the *pstSCAB* operon, and we also uncover metabolic deficiencies in the mutant strain – particularly its inability to thrive on L-lactate at concentrations approximating those in the chicken cecum - that likely contribute to loss of *in vivo* fitness.

## Results

### Mutagenesis of and characterization of the *C. jejuni pstSCAB* operon

We constructed an insertion allele using a cassette encoding resistance to kanamycin, inserted into the *pstS* gene of *C. jejuni* strain 81-176 (**Fig.S1**). The resulting kanamycin resistant strain was used in qRT-PCR experiments designed to analyze transcription of the *pstSCAB* locus. Consistent with the earlier RNASeq study (5), qRT-PCR of the wild type locus using primers specific for the junctions between open reading frames confirmed that the four genes are co-transcribed, a conclusion further supported by the observation that insertion in the *pstS* gene, the most upstream gene of the locus, resulted in loss of detectable downstream transcription (Fig. 1).

**Fig 1:**
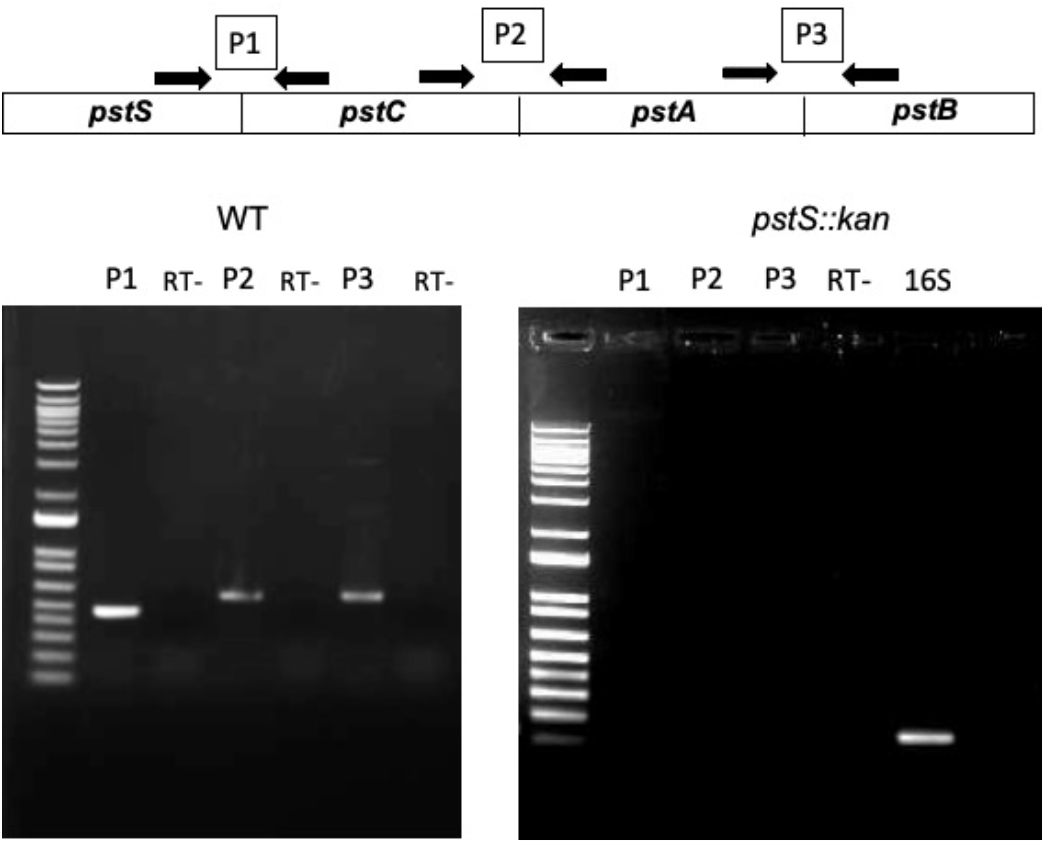
Characterization of *pstSCAB* operon: **A)** RT-PCR was performed to determine whether the four genes (*pstS*, *pstC*, *pstA* and *pstB*) are cotranscribed. Three intergenic primer sets (P1, P2 and P3) were designed to amplify transcripts crossing the gene boundaries. The product sizes in bp were correct as predicted in all cases. RT PCR with total RNAs were extracted from both DRH212 (left panel) and *pstS::kan* mutant (right panel) using the designed three primer sets. RT - negative control; 16S - as a positive control.

In the earlier RNASeq study abundance of the *pstS* mRNA was significantly higher than distal genes in the operon (5). Quantitative RT-PCR experiments confirmed that mRNA encoding *pstS* is more abundant than that of the *pstCAB* genes (**Fig. S2A)**. Differential stability within a polycistronic mRNA can be controlled through a conserved internal RNA hairpin that protects the transcript from RNA decay (14). The protective RNA structure is often present in the open reading frame of the downstream gene to protect the essential upstream gene from decay (14). Consistent with such a mechanism acting on the *pstSCAB* operon, we identified a thermodynamically stable hairpin loop structure (ΔG=-27.70) within the protein coding region of *pstC* gene (**Fig. S2B and C**). We hypothesize this RNA structure in *pstC* protects *pstS* mRNA from decay, leading to its elevated abundance relative to the other genes in the operon. Because of polarity from the *pstS::kan* insertion, for further experiments we used a complementing plasmid encoding the entire *pstSCAB* operon. In describing work with mutant and complemented strains, the strain is generally designated by its genotype (*pstS::kan*), and the strain with the mutation and carrying the complementing plasmid is designated *pstS::kan/C*.

### Phosphate storage and metabolism gene products are altered in the *pstSCAB* mutant

Survival of *C. jejuni* in stationary phase and nutrient limiting condition is dependent on polyphosphate (poly-P) metabolism (15). *C. jejuni ppk1* and *ppk2* encode poly-P kinases, with the former responsible for poly-P synthesis, using ATP generated from inorganic phosphate (16), and the latter generating GTP from poly-P (17). In some microbes Ppk2 can produce poly-P from GTP (17), but this activity has not been reported in *C. jejuni*. *C. jejuni* also encodes two exo-polyphosphatases, Ppx1 and Ppx2, which degrade poly-P into inorganic phosphate and have guanosine pentaphosphate hydrolase activity to generate guanosine tetraphosphate (ppGpp) (18). With the regulatory gene product SpoT, ppGpp regulates the stringent response of *C.jejuni*, contributing to survival in stationary phase (19). Another key gene of phosphate metabolism is *ppa*, which encodes an inorganic pyrophosphatase that hydrolyses pyrophosphate to inorganic phosphate (PPi→Pi)(19).

We investigated the effect of a *pstSCAB* mutation on these phosphate storage and stress response mechanisms, examining expression in wild type, mutant, and complemented mutant strains after 24 and 48 hours of growth in MHB. The gene largely responsible for poly-P synthesis, *ppk1*, was unchanged in its level of expression in the mutant versus the wild type at either time point. Expression of *ppk2*, which converts GDP to GTP using poly-P as a substrate - and is reported in some microbes (although not in *C. jejuni*) to produce poly-P from GTP - was elevated up to 10-fold in the mutant strain at 24 hours, returning to wild type expression levels by 48 hours (Fig. 2A). At 48 hours, expression levels of an exo-polyphosphatase gene (*ppx2*), the inorganic pyrophosphatase gene *ppa*, and the stringent regulatory gene *spoT* were significantly reduced in the mutant compared to wild type (Fig. 2B). As *ppx2* and *spoT* of *C.jejuni* regulate synthesis of ppGpp, important for survival in stationary phase, reduced expression of these two genes would likely result in decrease in the level of ppGpp in the *pstS::kan* mutant strain.

**Fig 2:**
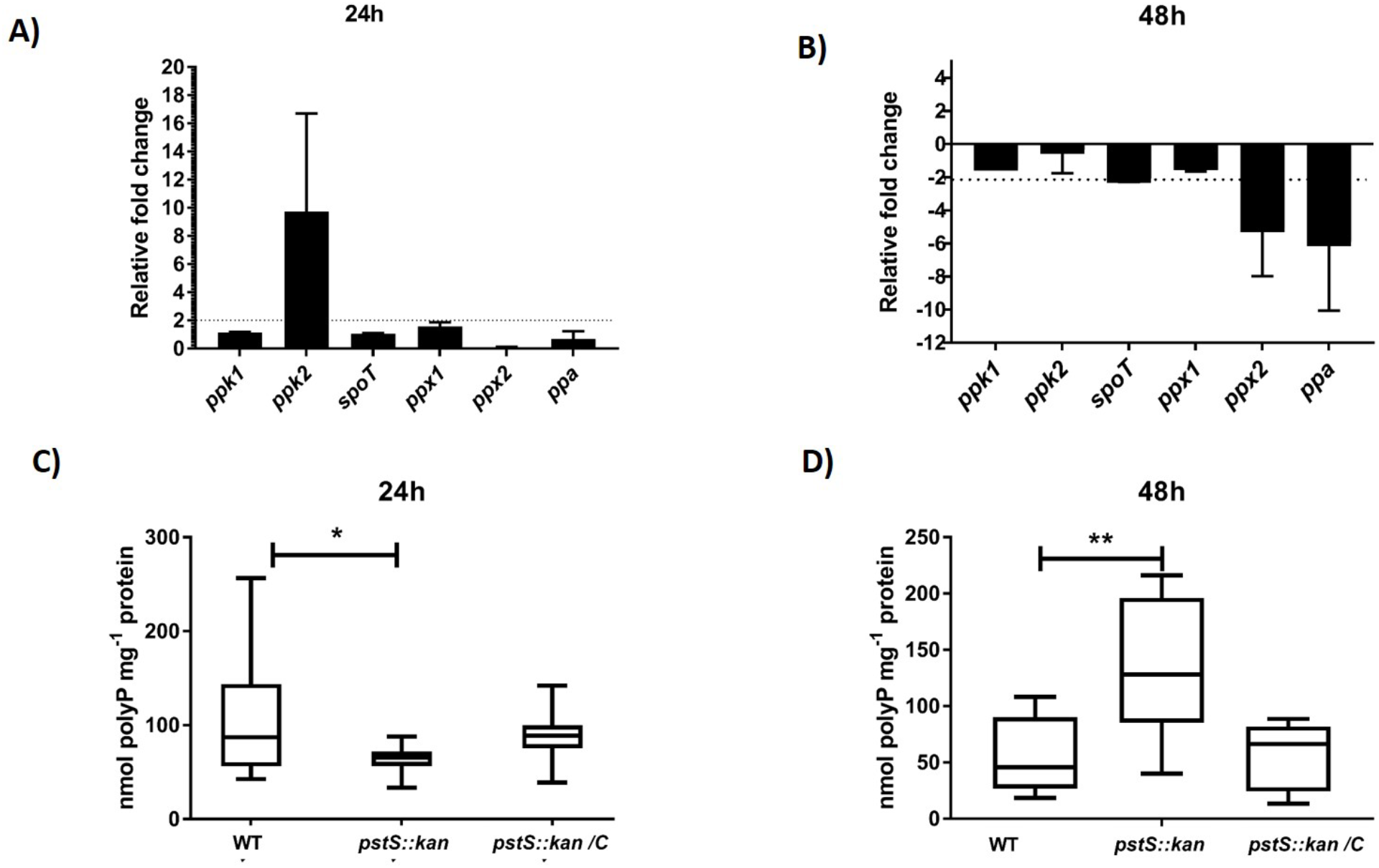
Comparative analysis of Poly-P metabolism regulatory genes and Poly-P level between DRH212 and *pstS::kan* mutant: **A)** and **B)** Transcription analysis of different stress regulatory genes *ppk1, ppk2, spoT* and *ppA* at 24h and 48h after growth of DRH212 and *pstS::kan* by Quantitative RT PCR. Changes of gene expression in the *pstS::kan* mutant compared with wild type DRH212 were determined by the 2^−∆∆CT^ method. Data are represented as mean value of three independent experiments with standard deviation (±). **C)** and **D)** Levels of Intracellular polyphosphate (Poly-P) of three strains (DRH212, *pstS::kan, pstS::kan/C*) were measured at two different time intervals 24h and 48h. Statistical analysis were done using Unpaired t-test and One way ANOVA with Holm-Sideak’s multiple test; (n=8,±SD), ∗, P<0.05; ∗∗, *P* < 0.005.

At 24 hours, poly-P levels in the mutant were roughly half of those observed in the wild type. At 48 hours, by which time the mutant had diminished in viability by several orders of magnitude (see below, Fig. 3B), poly-P was elevated to about three times the levels observed in wild type (Fig. 2C and D). With elevated *ppk2* expression at 24 hours in the *pstS::kan* mutant, decreased intracellular poly-P might be expected, as the Ppk2 enzyme consumes poly-P to convert GDP to GTP. The higher levels of poly-P observed in the *pstS::kan* mutant compared to wild type at 48 hours is readily explained by decreased expression of the exopolyphosphatase gene *ppx2* in the mutant background, and is consistent with a similar report of a *pstS::kan* mutant of *Pseudomonas aeruginosa* (20).

**Fig 3:**
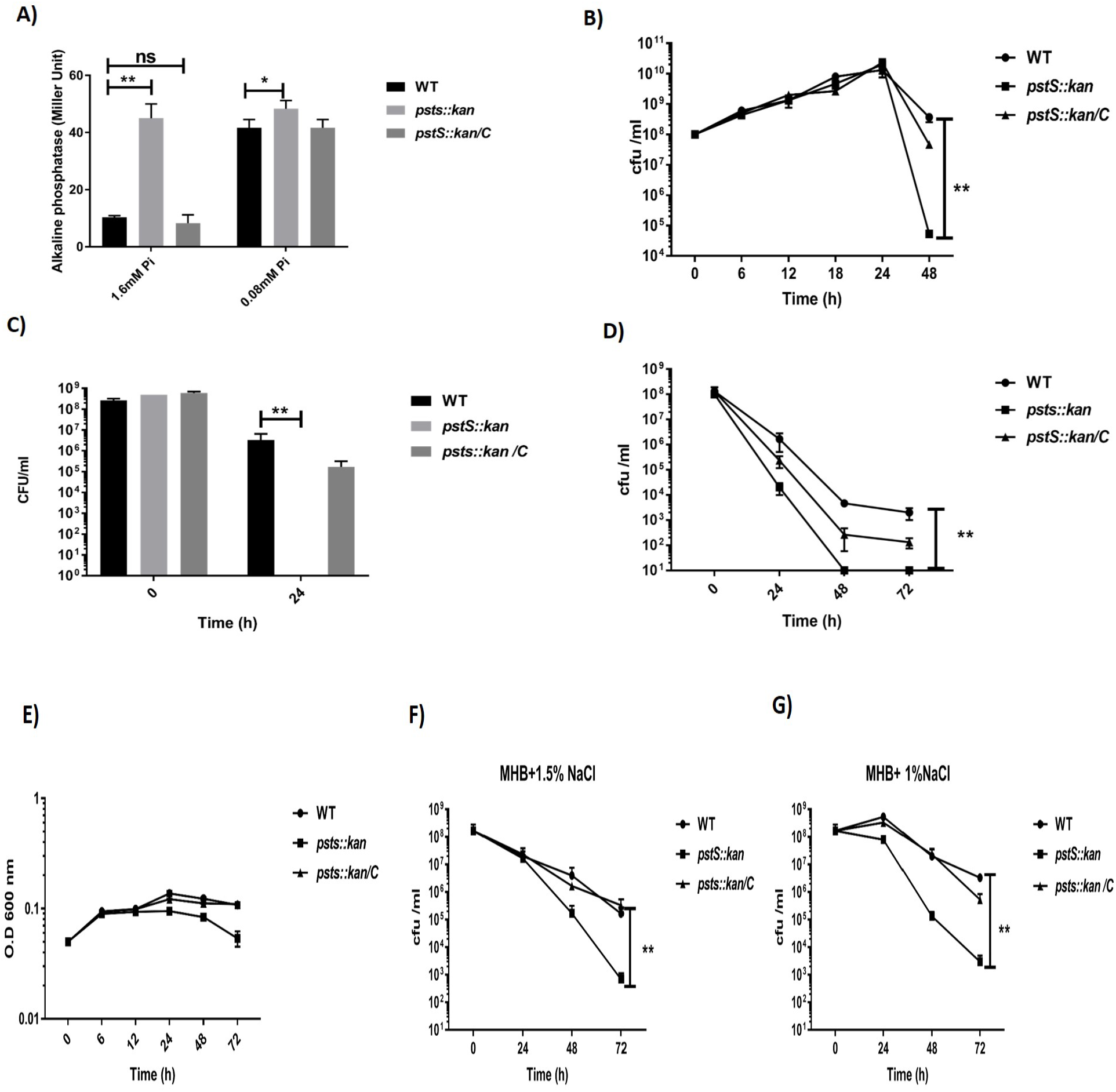
Characterization of *C. jejuni pstS::kan* strain in different *in vitro* conditions: **A)** Alkaline phosphatase activity was determined in pyruvate-minimal media with 1.6 mM or 0.08 mM Pi (n=3). Statistical analysis was done using two way ANOVA with Sidak’s multiple comparisons test. ∗ **,P<0.05;** ∗∗, ***P* < 0.005**, **B)** *C. jejuni* strains were grown in Mueller-Hilton (MH) broth in microaerobic condition for 48 h and after different time intervals CFU/ml were determined**. C)** *C. jejuni* DRH212 was grown in MH broth for 48h. After 48h, sterile spent media were collected by filter sterilization. Mid-log phase cultures of DRH212, *pstS::kan, pstS::kan/C* strains were inoculated (10^8^ CFU/ml) in spent media and survival was measured at different time intervals by CFU count. **D)** In nutrient down shift assay, mid-log phase of DRH212, *pstS::kan, pstS::kan/C* strains were inoculated in minimal media where no carbon and phosphate source were present. The survival assay was measured by CFU count at different time intervals. **E)** Osmotic stress tolerance: strains were inoculated in MH broth containing 1% NaCl (**F**) or 1.5% NaCl (**G**) and growth was determined by measuring O.D at 600nm and cfu counts. Each experiment represents mean value of three independent experiments. Statistical significant was done student t test. ∗, P<0.05; ∗∗, *P* < 0.005.

### *pstS* is essential for stationary phase survival and osmotic stress response

Expression of *pstS* is elevated approximately four-hundred-fold in 0.08 mM inorganic phosphate (Pi) compared with 1.6 mM Pi, through activation of the PhoRS system at the more limiting phosphate concentration (7). The PhoRS-regulated alkaline phosphatase gene of *C. jejuni* (*phoX*) is also induced at lower phosphate concentration, providing Pi for transport into the cell by the PstSCAB system (21). We hypothesized that the *pstS::kan* mutant strain would signal phosphate limitation to the cell, even at a concentration (1.6 mM) that does not typically result in *phoX* expression. We measured alkaline phosphatase activity in wild type and *pstS::kan* mutant cells grown in 0.08 mM and 1.6 mM phosphate. Supporting our hypothesis, the mutant expressed significantly elevated alkaline phosphatase activity at both the lower (0.08 mM) and higher (1.6 mM) concentrations, whereas wild type (and the complemented mutant) expressed high levels of alkaline phosphatase activity only at the lower phosphate concentration (Fig. 3A). We conclude that, even at elevated phosphate concentrations, the *pstS::kan* mutant strain senses phosphate limitation.

To assess the effect of loss of *pstSCAB* on growth of *C. jejuni*, we cultured wild type, mutant, and complemented mutant strains in Mueller-Hinton broth (MHB), a complex medium of protein hydrolysate and infusion. All grew to similar levels for 24 hours, peaking at 10^10^ cfu/ml (Fig. 3B). By 48 hours, the wild type and complemented mutant strains had dropped about two orders of magnitude in plating efficiency, while the mutant strain had dropped by over five orders of magnitude in plating efficiency, suggesting a severe defect in survival in stationary phase and consistent with the diminished levels of poly-P we detected in the cells at that time-point. Assessing survivability in media depleted of nutrients provided a more direct examination of the challenge posed by loss of *pstSCAB*. In this experiment the wild type strain was grown for 48h in MH broth to reach stationary phase, after which the growth media was collected by filter sterilization. We inoculated mid-log phase wild type, *pstS::kan* and the complemented mutant in the sterilized depleted media, with initial inoculum (5×10^8^ CFU/ml), and assessed viable cells at 24 hours. In contrast to wild type and complemented mutant, which were still recoverable at high numbers in depleted media, the mutant was completely unrecoverable (Fig. 3C) confirming that loss of the phosphate uptake system causes a significant defect in growth physiology. Likely contributing to this loss of fitness under limiting nutrient conditions (Fig. 3D) is the decreased expression demonstrated (Fig. 2B) in of *ppx2* and *spoT*, regulators of ppGpp synthesis.

In addition to stationary phase survival, poly-P metabolism contributes to survival under osmotic stress, based on analysis of strains carrying mutations in the genes described above (16). To assess whether phosphate transport *per se* contributes to osmotic survival, we analyzed wild type and *pstS::kan* mutant growth and survival at 1% and 1.5% NaCl, which approximate concentrations within the chicken gastrointestinal tract (22). The *pstS::kan* mutant strain demonstrated a significant growth defect (Fig. 3E) and lower plating efficiency than wild type on 1% and 1.5% NaCl (Fig. 3F and G). The mutant strain was restored to wild type levels of plating by complementing with *pstSCAB* in trans, confirming that loss of viability at elevated osmolarity is due to loss of phosphate transport. Thus, similar to mutant strains lacking *ppx1* and *ppx2* (exo-polyphosphatases) (18), which are unable to degrade polyphosphate, the phosphate transporter mutant survives poorly under osmotic stress conditions that are similar to those found in the chicken gastrointestinal tract.

### Loss of PstSCAB compromises *C. jejuni* chicken colonization and fitness

We investigated whether elevated abundance of *pstS* mRNA in *C. jejuni* harvested from chick ceca, compared to *C. jejuni* grown *in vitro* (5), correlates to actual requirement for *pstSCAB* during growth in the chicken. We orally inoculated day-of-hatch chickens with two different doses (10^3^ or 10^6^ CFU) of wild type *C. jejuni* 81-176, the *pstS::kan* mutant derivative, and the complemented mutant, and enumerated *C. jejuni* loads in cecal contents on days three and seven after inoculation. Irrespective of the inoculum dose, the mutant strain exhibited a large colonization defect at both days three and seven compared with wild type (Fig 4A,B,C and D), being recovered approximately five orders of magnitude less than wild type when introduced at the lower inoculum and by roughly two orders of magnitude when introduced at the higher inoculum. In all cases the complemented strain colonized at levels similar to wild type.

**Fig 4:**
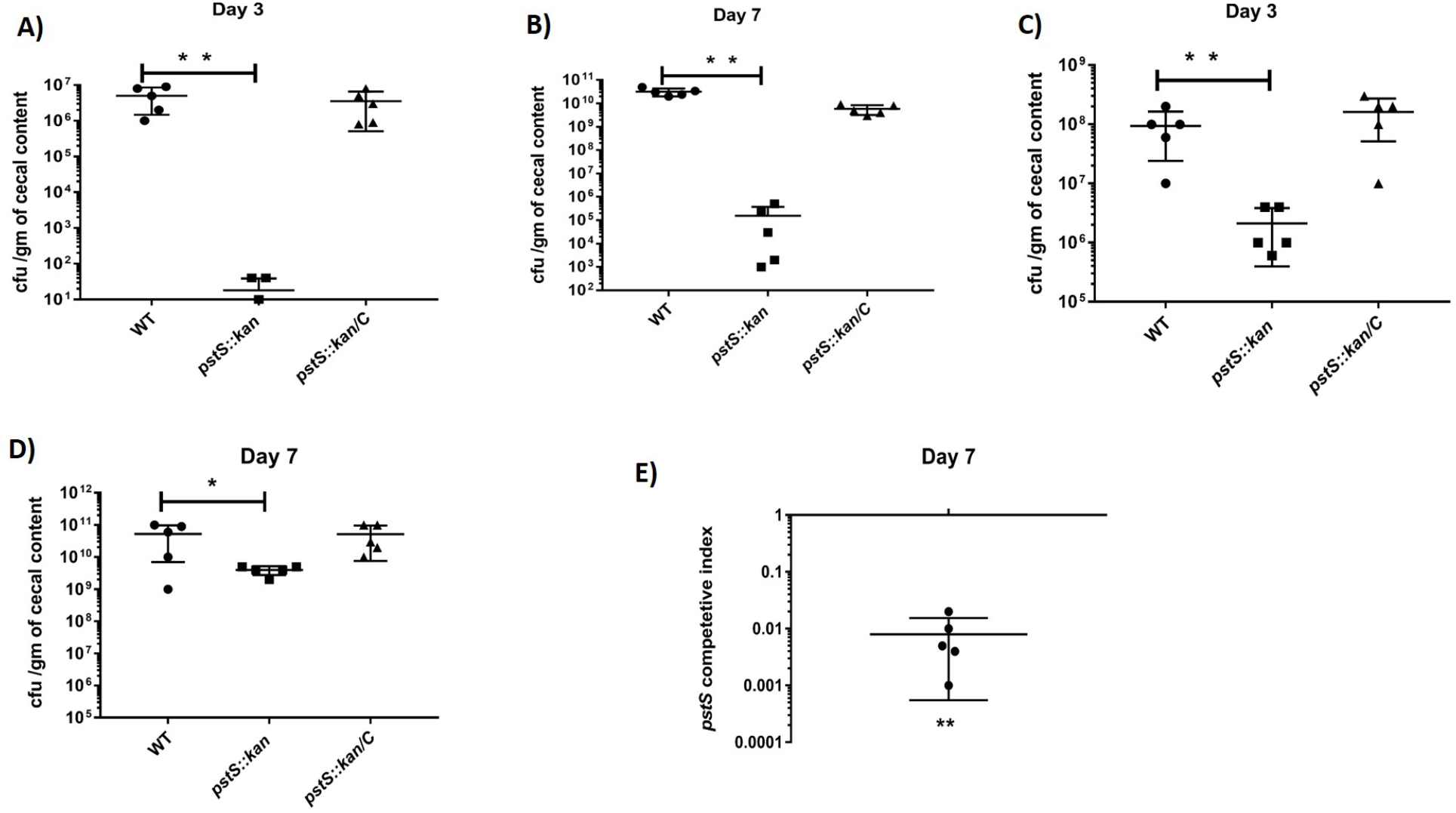
Colonization ability of *pstS::kan* mutant in chickens: Day-of-hatch white Leghorn chicks were infected with either DRH212, *pstS::kan* or *pstS::kan/C* strains of *C. jejuni* (two different doses 10^3^ and 10^6^ CFU/ml) and cecal loads were enumerated at day three and day seven. Comparative colonization ability between DRH212 and *pstS::kan* strains at day three and day seven after oral inoculation with 10^3^ (**A, B)** and 10^6^ CFU/ml **(C, D).** Statistical analysis was one way ANOVA with Tukey’s multiple comparison test (n=5,±SD). **E)** In competition analysis, DRH212 versus *pstS::kan* mutant, day-of-hatch chicks were infected with equal ratio of DRH212 and *pstS::kan* mutant. At day seven the ratio of *pstS::kan* mutant to DRH212 was determined and is presented as a competitive index. Statistical analysis was performed using a one-simple t test against a hypothetical value of 1. (n=5 of each group). ∗, P<0.05; ∗∗, *P* < 0.005.

In addition to individual colonization assays with wild type and *pstSCAB* mutant strains, we carried out a competition assay between the wild type strain 81-176 and the *pstSCAB* derivative, mixing the strains at 1:1 concentration in the inoculum and introducing the combination at 10^6^ CFU/ml by oral gavage to day-of-hatch chickens. We harvested cecal contents on day seven post-infection and determined the ratios of the mutant and wild type strains by measuring relative numbers of kanamycin resistant (*pstSCAB* mutant) and sensitive (wild type) *C. jejuni* in the output (Fig 4E). The wild type strain exhibited greater fitness in this experiment than the mutant by 100-fold. The wild type strain also out-competed the mutant when grown together in nutrient-rich media or in minimal media supplemented with 80 uM phosphate (**Fig. S4A and B)**. These findings indicate the essential nature of *pstSCAB* for chick colonization and extend the earlier observation of high-level *in vivo* expression of this operon in the transcriptome study (5).

### The phosphate transporter is essential for lactate-dependent growth

Lactate is present in both D- and L-isomers at varying concentrations in the chicken gastrointestinal tract, generally decreasing from the upper to the lower intestine and ceca (23). Although at high concentrations lactate inhibits growth of *C. jejuni* (24) and reduces expression of some colonization traits (23), many strains - including 81-176 - have pathways for uptake of and growth in lactate, including a transporter for L-lactate encoded by *lctP* (CJJ81176_0113) and two NAD-independent membrane-associated L-lactate dehydrogenase complexes (CJJ81176_0112-0110) and (CJJ81176_1182), which convert L-lactate to pyruvate (25). Mechanisms for uptake and assimilation of D-lactate are less understood (25).

Given the reduced fitness of the *pstSCAB* mutant in the cecum, we investigated the growth of the mutant on this potentially relevant carbon source in that environment. We cultured wild type, mutant, and complemented mutant strains on minimal media containing D- + L-lactate (10 mM total lactate) in both low (80 micromolar) and high (1.6 millimolar) phosphate concentrations. In both phosphate concentrations, the mutant grew to levels 100-1000 times lower than wild type at 24 and 48 hours of growth (Fig. 5A and B). This contrasts to cells cultured in media with pyruvate as the carbon source, where mutant and wild type grew to similar levels in high phosphate media (1.6 mM) and differed by only an order of magnitude at the lower phosphate concentration (80 µM; **Fig. S5A-B**). When cultured on L-lactate alone, the *pstS::kan* mutant strain was significantly limited compared to wild type, irrespective of phosphate concentration, while on D-lactate alone, the mutant was diminished for growth during initial exponential growth phase (irrespective of phosphate levels) but ultimately attained the same level of growth as the wild type and the complemented mutant (Fig. 6A-D). Collectively these data suggest that the phosphate transport mutant is challenged for growth on L-lactate in particular; even when both D- and L-lactate are available *in vitro* the mutant is at a significant disadvantage compared to wild type (Fig. 5). Given that D- and L-lactate may serve as growth substrates for *C. jejuni* growth in the chick cecum (25), these data also contribute to explaining the fitness disadvantage of the mutant *in vivo*.

**Fig 5:**
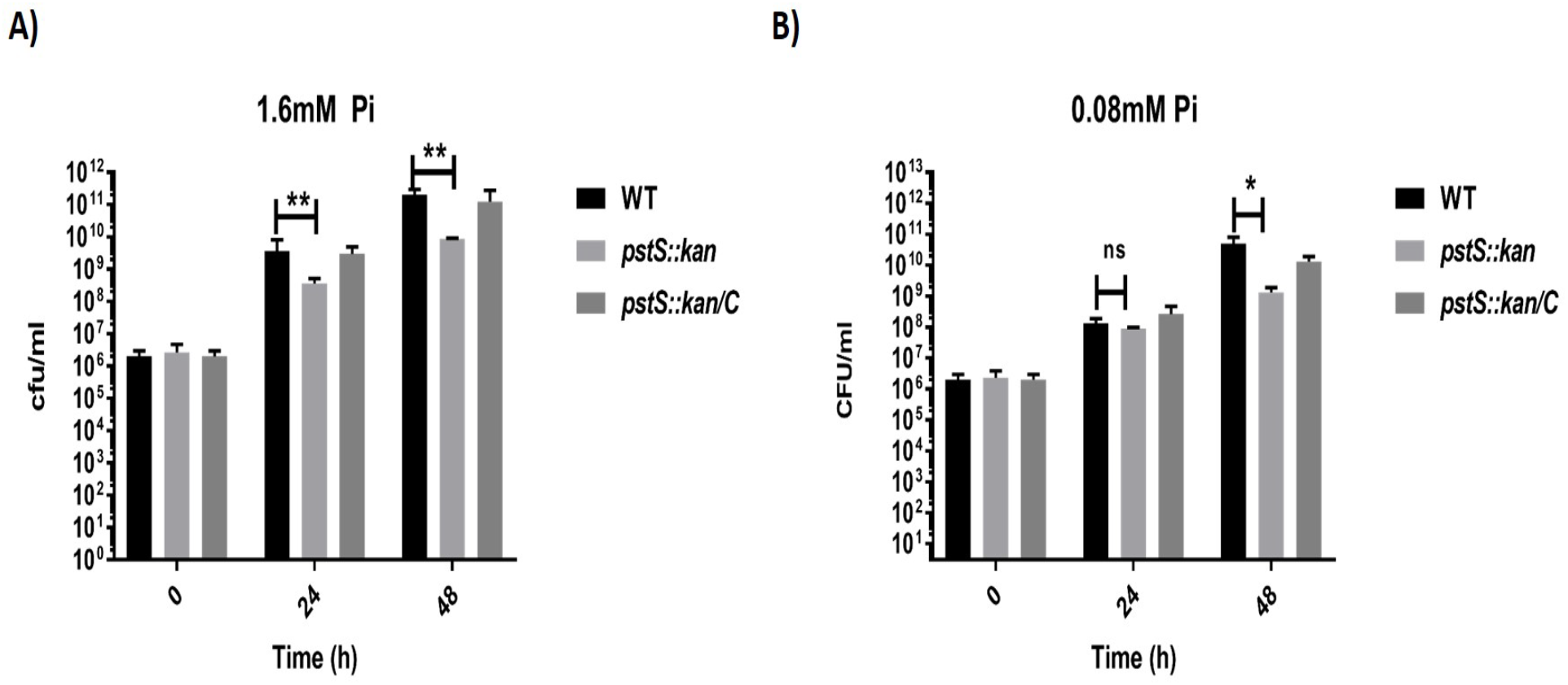
PstSCAB transporter of *C. jejuni* is essential for growth in lactate: *C. jejuni* strains were grown in minimal media containing both D+L lactate (each 5mM) with two concentrations of inorganic phosphate. Growth of DRH212 and *pstS::kan* was measured by CFU count in minimal media (D+L lactate) with **A)** 1.6mM Pi and **B)** 0.08mM Pi at 24 and 48h. N=3; error bar indicated SD. Statistical analysis was done by two-way analysis of variation (ANOVAs) with Sidak’s multiple comparisions test. ∗, P<0.05; ∗∗, *P* < 0.005.

**Fig 6:**
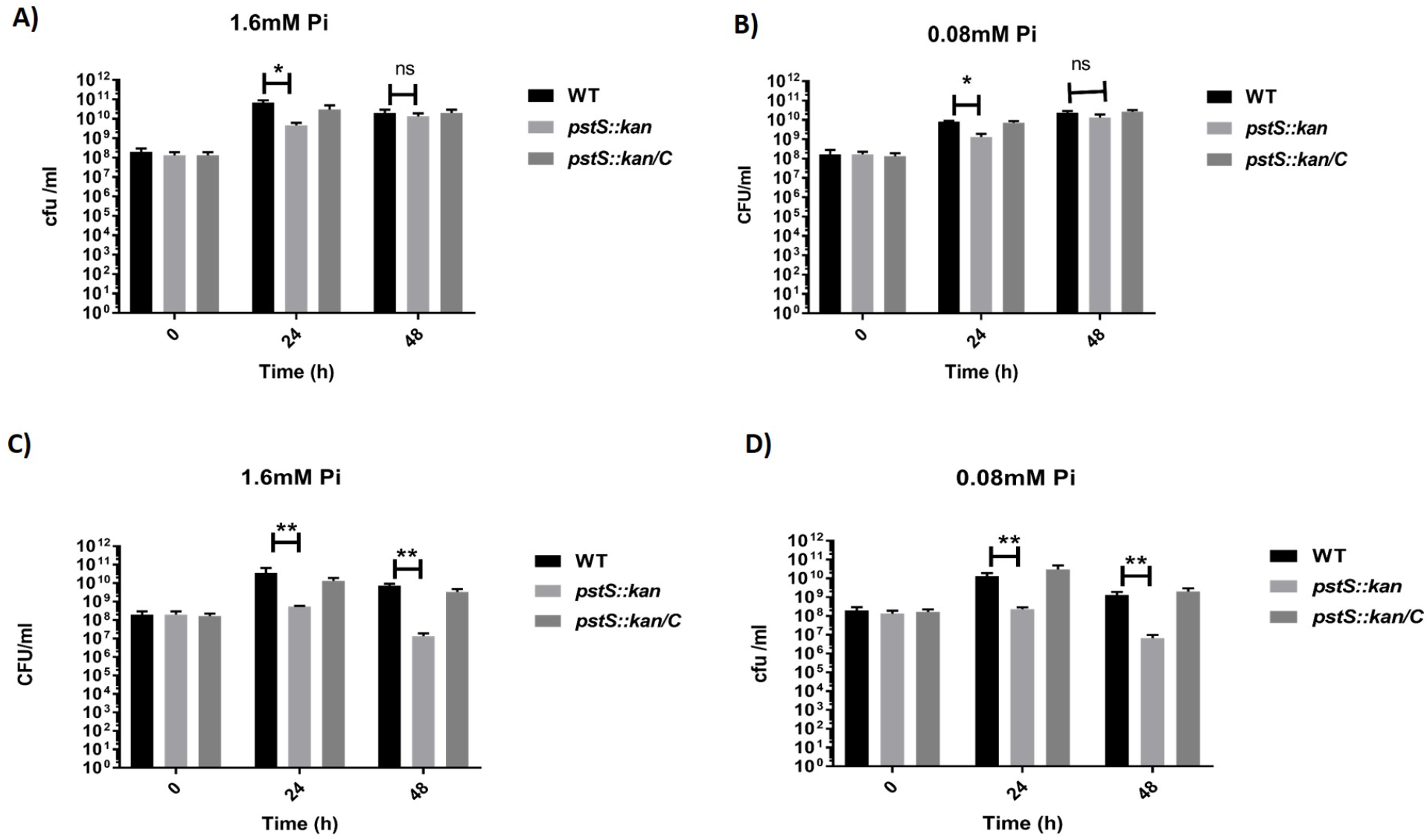
Comparative growth analysis of *C. jejuni* strains in minimal media with D- or L-lactate. *C. jejuni* strains were grown in minimal media with D-lactate or L-lactate as a carbon source and high or low concentration of inorganic phosphate. Growth of *C. jejuni* strains was measured in D-lactate minimal medium with **A)** 1.6mM and **B)** 0.08mM phosphate by CFU counts at 24h and 48h. Growth of these three strains was also determined in L-lactate minimal medium with 1.6mM **C)** and 0.08mM **D)** phosphate. N=3; error bar indicated SD. Statistical analysis was done by two-way analysis of variation (ANOVAs) with Sidak’s multiple comparisons test. ∗, P<0.05; ∗∗, *P* < 0.005.

Lactate-derived pyruvate is converted to acetate in *C. jejuni* via phosphotransacetylase (Pta) and acetate kinase (AckA); this acetogenesis pathway is critical for establishing chicken colonization (23). Genes controlled by this acetogenesis pathway may contribute to fitness in the chick ceca including those encoding γ-glutamyl transferase (Ggt), a component of an amino acid transporter complex (Peb1c), and a putative di/tri-peptide transporter (Cjj81176_0683) (23). The higher L-lactate levels in the upper intestine, compared to those of the lower intestine and ceca, lead to reduced expression of these genes through an undefined regulatory mechanism that correlates to relatively lower levels of *C. jejuni* in the upper intestine (23). We hypothesized that the poor growth on lactate of the *pstSCAB* mutant might also correlate with diminished expression of acetogenesis-regulated colonization traits. To test this, we carried out quantitative RT-PCR to measure transcript levels of *ggt*, *peb1c* and *Cjj81176_0683*. Expression of each was reduced by two-to-four-fold in the mutant compared to wild type (Fig 7). This reduction in expression is similar in magnitude to their reduced expression in the upper intestine where *C. jejuni* colonizes poorly (23). Thus, we conclude that in addition to poor growth on L-lactate by the mutant lacking PstSCAB, the strain also expresses potentially key colonization determinants at levels apparently insufficient for cecal colonization.

**Fig 7:**
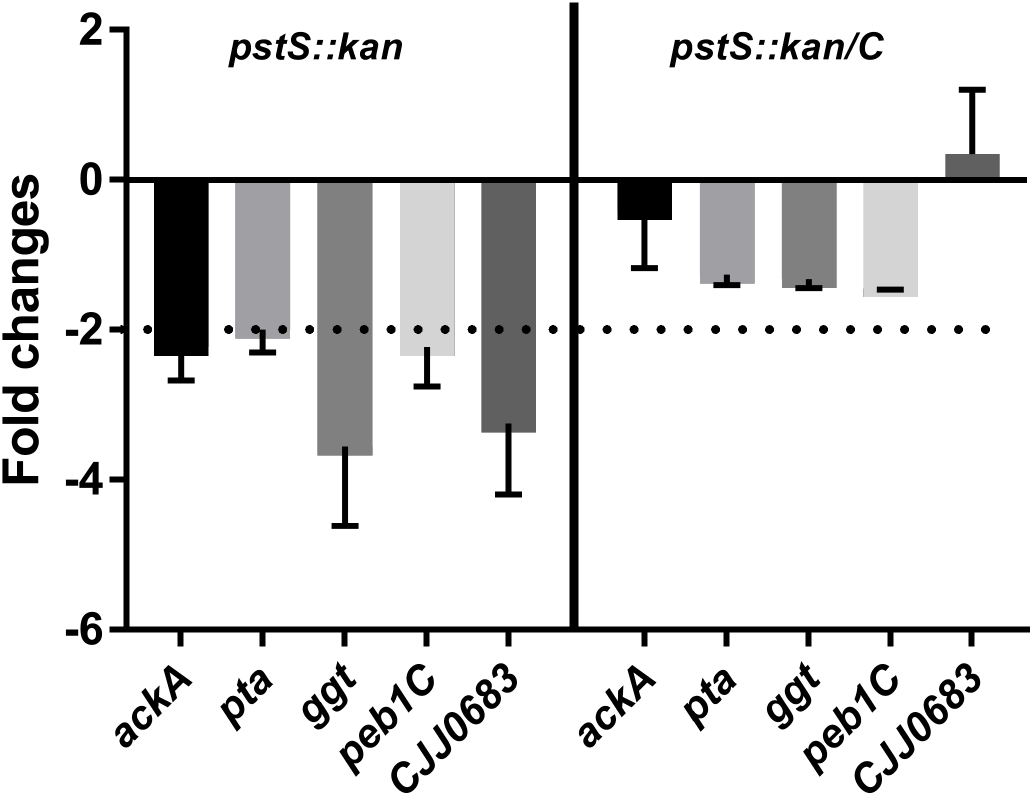
Transcript analysis of acetogenesis regulatory and acetogenesis dependent gene expression in wild type, *pstS::kan*, and complemented strains: Bacterial strains were grown in minimal media containing L-lactate with 1.6mM Pi up to 18h, after which cells were harvested for RNA isolation. Acetogenesis regulatory genes (*ackA* and *pta*) and acetogenesis dependent genes (*ggt,peb1C* and *CJJ0683*) were measured in *pstS::kan* mutant and *pstS::kan/C* strains relative to DRH212 by real time PCR. N=3; error bar indicated SD.

### Metabolomic analysis of wild type and *pstSCAB* mutant *C. jejuni*

To explore further the growth physiology of wild type and *pstSCAB* mutant on lactate, we measured intracellular metabolites using mass spectrometry. We observed no significant difference in intracellular lactate or pyruvate between wild type and the *pstS::kan* mutant strain (**Fig S6A)** indicating that the phosphate transport system has no effect on lactate uptake or its conversion to pyruvate. Of metabolites we measured, adenine - and its deamination product hypoxanthine - were significantly lower in the mutant compared with the wild type (Fig. 8 B and C), and this was reflected in diminished ATP levels in the mutant as well. (Fig. 8 A). The low intracellular Pi concentration in the *pstS::kan* mutant (as reflected in elevated alkaline phosphatase activity; Fig. 3A), correlated to decreased intracellular ATP levels and might account in part for its poor fitness. This was confirmed with ATP assays on extracts of cells cultured for 24 hours in minimal-L-lactate-media demonstrating intracellular ATP concentrations in the *pstS::kan* mutant significantly below those of wild type, irrespective of phosphate concentration (Fig 8A). *C. jejuni* does not encode a purine/pyrimidine or nucleotide/nucleoside transporter (26), and indeed adenine supplementation (1uM or 10uM) of L-lactate media did not restore growth or ATP levels in the mutant (**Fig. S7**).

**Fig 8:**
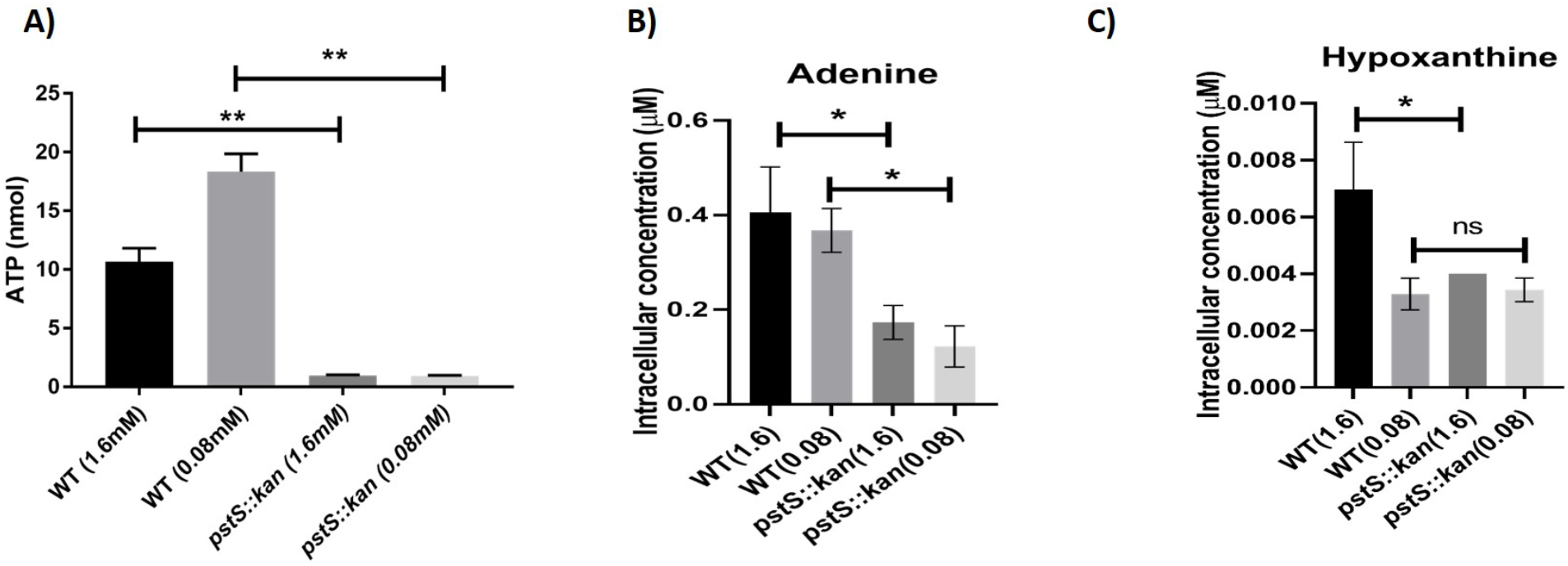
Intracellular ATP and metabolites in DRH212 and *pstS::kan* mutant grown in L-lactate minimal media. DRH212 and *pstS::kan* strains were grown in L-lactate minimal media with 1.6mM and 0.08mM Pi at 37^0^ C. After 18h, **A)** ATP levels in DRH212, *pstS::kan* were detetermined. N=3; error bar indicated SD. The intracellular concentration of **B)** adenine and **C)** hypoxanthine were measured by liquid chromatography mass spectroscopy analysis. Data represent mean value of two independent experiments. The statistical analysis was done by one way ANOVA. ∗, P<0.05; ∗∗, *P* < 0.005.

## Discussion

The *pstSCAB* phosphate transporter locus of *C. jejuni* is among the more significantly expressed transcripts identified by RNASeq from microbes harvested from the chick cecum (5) and during infection of humans (27). A transcriptomic study of *C. jejuni* colonizing chicks using microarray analyses (28) did not report high level expression of *pstSCAB*, although there are significant differences in the way the RNASeq and microarray studies were carried out with regards to chicken inoculation and colonization. The RNASeq study (5) used an inoculum of 10^3^ CFU of *C. jejuni*, and harvested cecal contents for RNA preparation after seven days of colonization. In contrast the microarray study used a much higher inoculum of 10^9^ CFU and harvested cecal contents after only 12 hours of colonization (28). It is conceivable that these differences in inocula and time of colonization would result in different physiological outcomes in the *C. jejuni* recovered from the cecal contents.

Generally, elevated level of expression of the high-affinity phosphate transporter in bacteria is observed in a low-phosphate environment. For example, transcription of *pstS* in *C.jejun*i is significantly increased at 0.08 mM concentration compared to 1.6 mM (7). We determined phosphate concentrations within the chicken cecum as relatively high, 0.5-0.8 mM/100mg of cecal content (**Fig. S3)**. Further, previous metabolite studies estimated that the Pi concentration in intestinal lumen of healthy adults are varied between 15-30 mM on a Western diet (29). At these *in vivo* levels of inorganic phosphate, we would not expect elevated expression of *pstSCAB* during colonization in chicks and humans, so it remains unclear what drives this outcome during host colonization. We hypothesize that, although present at high detectable levels, phosphate is nevertheless unavailable to *C. jejuni*, perhaps due to competition from other microbes. We previously demonstrated *in vivo* competition for zinc, resulting in poor colonization of a *C. jejuni* mutant lacking the *znuA* high-affinity zinc transporter. In that case, the colonization defect of the *znuA* mutant was mitigated in chicks reared in germ-free conditions harboring a limited microbiota (30).

The *pstSCAB* mutant of *C. jejuni* showed a significant colonization defect on chicken at both inocula we tested (10^3^ and 10^6^ CFU), with the most significant defect at day three after infection. From these data, we hypothesize that *pstSCAB* enables *C. jejuni* to survive initial conditions after reaching the chicken gut. We also observed a dose dependent colonization defect of the mutant at day seven. A previous study reported similar colonization levels between wild type *C. jejuni* and a mutant lacking *phoRS*, which encodes a regulator of *pstSCAB* expression (6). That study used a different strain of *C. jejuni*, and the animal data were not shown, so we do not know what infection dose was used or what time points were assessed post-inoculation. Another study demonstrated that mutants lacking the alkaline phosphatase PhoX have diminished capacity for colonizing chickens compared to wild type (21) and that this is a dose-dependent outcome with somewhat greater colonization defect of the mutant observed after a low dose (10^3^) inoculum by comparison to a higher dose inoculum (10^5^).

As a proxy for analyzing fitness of wild type and *pstSCAB* mutant in humans, we measured growth of WT and *pstSCAB* mutant strain *in vitro* in minimal media with 10% human fecal extract, in which the phosphate concentration was approximately 500 µM (**Fig. S8A)**. The mutant strain grew in this medium after 24 hours, but nevertheless grew to levels 10x lower than wild type and the complemented mutant (**Fig. S8B**). Taken together with previous human transcriptomic data demonstrating elevated levels of *pst* messages during human infection (27), we conclude that phosphate transporter of *C. jejuni* is required for fitness in both chickens and humans.

The *pstSCAB* operon is transcribed as a full-length polycistronic mRNA, demonstrated both in the earlier RNASeq study and in our qRT-PCR presented in this work. Trancripts encoded by the first gene in this operon, *pstS*, are present at much higher levels than those of the three downstream genes (5). We postulate that a potential stem-loop structure encoded just downstream of the *pstS* gene, which we postulate serves to stabilize *pstS* mRNA from degradation, resulting in higher steady state levels compared to the other genes of the operon. A similar mechanism is proposed for the *pstS2* operon of *Vibrio cholerae* (31).

In complex media, PstSCAB is dispensable until late in growth, after which it is required for cells to remain viable in the nutrient-depleted environment (Fig.3). As transcription of the locus is elevated during mid-logarithmic growth (compared with stationary phase growth (5)) we conclude that assembly of the system and Pi uptake occurs early during growth and enables survival later when the locus is far less transcriptionally active. Accumulation of poly-P in the *pstSCAB* mutant late in growth may be the result of decreased expression of the exo-polyphosphatase genes *ppx1* and *ppx2*, which encode enzymes that metabolize poly-P. Consistent with a need to balance poly-P production and metabolism for wild type fitness in *C. jejuni* others, have demonstrated that *phoX* and *ppk1/ppk2* mutants also have survival defects in nutrient limiting condition (18). Increased poly-P was also observed in phosphate transport mutants of *M. tuberculosis* (32).

The failure of the *pstSCAB* mutant to thrive - and its loss of viability - in 10 mM lactate, on which wild type grows to high levels, is worth further analysis. The mutant strain exhibits reduced expression of the *pta-ackA* genes in the acetogenesis pathway, which could be partly responsible for the decreased ATP levels in the mutant compared to the wild type. Lactate at high concentrations causes membrane damage to *C. jejuni* and is the basis for the suggestion that *Lactobacillus* strains that produce high lactic acid levels might be useful probiotics to reduce *C. jejuni* loads in poultry (24). Concentrations required for killing *C. jejuni* are much higher than 10 mM, but in the context of loss of *pstSCAB* perhaps the cell is sensitized to lactate toxicity. A mutant of avian pathogenic *E. coli* lacking *pst* was demonstrated to be more sensitive to membrane-targeting polymyxin B, but we did not observe that with *pstSCAB* mutant *C. jejuni*. How *pstSCAB* contributes to lactate-dependent growth will be the subject of future research.

## Methods and Materials

### Bacterial Strains and Media

All strains and plasmids used in the study are listed in Table 1. The wild type strain used in all experiments is DRH212, a streptomycin resistant derivative of *C. jejuni* 81-176 (33). All *C. jejuni* strains were routinely grown under microaerobic conditions (85% N2,10%CO2, 5% O2) on Muller-Hilton (MH) agar or in broth under necessary antibiotics at the following concentrations: chloramphenicol, 15 µg ml^−1^; kanamycin, 50 µg ml^−1^; TMP, 10 µg ml^−1^; streptomycin, 2 mg ml^−1^. *C. jejuni* strains were stored in MH broth with 20% glycerol at −80°C.

**Table 1:**

| Strain or plasmid | Relevant characteristics | Source of Reference |
| --- | --- | --- |
| *C. jejuni* DRH212 | Streptomycin-resistant wild type *C. jejuni* 81-176 (Sm<sup>r</sup>) | 34 |
| DRH212 *pstS::kan* | *pstS* insertion mutant of *C. jejuni* DRH212 (Kn<sup>r</sup> Sm<sup>r</sup>) | In this study |
| DRH212 *pstS::kan/C* | *pstS* insertion mutant of *C. jejuni* DRH212 content *pstSCAB* cloned PECO212 ((Kn<sup>r</sup> Sm<sup>r</sup> Cm<sup>r</sup>) | In this study |
| *E.coli* DH5a | *E. coli* strain used for cloning |  |
| *E.coli* DH5a/pRK212.1 | Contains conjugative plasmid for the conjugation of plasmid DNA into *Campylobacter* | 36 |
| Plasmids |  |  |
| pGEMT | Subcloning vector (Ap<sup>r</sup>) |  |
| pECO102 | pRY112 derivative with *cat* promoter in XhoI-BamHI | 36 |
| pILL600 | contains *Campylobacter* kanamycin cassette | 36 |

### Construction of a *pstS::kan* insertion mutant and complementation

*C. jejuni pstS*::*kan* mutant was constructed by insertion of kanamycin resistance cassette into the gene as previously described (34). Briefly, the 500bp of both upstream and downstream regions of *pstS* locus were amplified by PCR with flanking region containing SmaI restriction site. The amplified upstream and downstream region were joined by Soeing PCR. The 1000bp amplified PCR products was cloned into pGEM-T easy. The *Campylobacter* kanamycin resistance cassette was excised from PILL600 using SmaI and ligated between upstream and downstream cloned region of pGEM-T construct. The plasmid was electroporated into DRH212 strain and grown on MH agar overnight at 37°C under microaerobic conditions. Then cells were plated and grown on MH agar containing kanamycin for 2-3 days, and successful integration of *pstS::kan* into the chromosome was confirmed by PCR. As we demonstrate in Results, this insertion is polar on the *pst* genes downstream of *pstS*. Therefore, for complementation, the coding sequences of whole *pstSCAB* operon - from the second codon of *pstS* to the stop codon of *pstB* - were amplified by PCR and cloned into the BamHI site of pECO102 plasmid (35), which was introduced into *E. coli*DH5α/pRK212.1 strain (35). Conjugation to *C. jejuni* was performed as previously described by Guerry et al (36).

### Growth Curve analysis in MH broth and Minimal media

*C. jejuni* strains WT, *pstS::kan*, *pstS::kan/C* were grown from freezer stocks on MH agar containing desired antibiotics for 24h under microaerobic condition. The strains again re-streaked and grown for 16h. The bacteria were suspended in MH broth and inoculated in 25ml of MH broth containing desired antibiotics. The initial O.D was approximately 0.05 and bacterial numbers were measured at different time intervals up to 48h. To determine growth of *C. jejuni* strains in different concentrations of inorganic phosphate, strains were grown in MCLMAN minimal media containing different carbon sources including sodium L-pyruvate, D-lactate and L-lactate (37) as noted. Strains were washed twice with minimal media and inoculated with 25ml of minimal containing different concentration of Pi. Then, the growth was measured via absorbance at 600nm (OD_600_) up to 48h with the initial O.D was 0.05.

### ATP measurements

ATP levels in strains were determined by BacTitre-Glo^TM^ Assay Kit (Promega) as previous described. Briefly, strains were grown for 24h in minimal media (L-lactate) with high (1.6mM) and lower concentration (0.08mM) of Pi and normalized OD at 0.3. Then ATP level were measured according to the manufacture’s protocol and ATP concentrations were measure by standard curve (26).

### Measurement of intracellular metabolites

Wild type strain DRH212, *pstS::kan* were grown in presence of L-lactate with higher and lower concentration of phosphate for 18h. The cells were collected by centrifugation and washed twice with normal saline. Then all samples were normalized approximately with OD. 1. Then the samples were extracted for determination of metabolites. Samples on dry ice were combined with 800 microliters of ice-cold acetonitrile:methanol:acetone (8:1:1, v:v:v) and suitable stable isotope-labeled or synthetic internal standards including D4-glycochenodeoxycholic acid and myristoyl phosphatidylcholine for estimation of metabolite recovery and for relative quantitation across experimental groups. Samples were homogenized with zirconium oxide beads for 2 minutes in a Bullet Blender (Next Advance) at 4 degrees Celsius, incubated for 30 minutes at 4 degrees Celsius, then centrifuged for ten minutes at 10,000g at 4 degrees Celsius. Supernatants were filtered through 0.2 micron syringe filters (Fisher Scientific), and evaporated to dryness in a SpeedVac. Samples were resuspended in 95% acetonitrile for use in LC-MS analysis.

### Liquid chromatography-mass spectrometry

Untargeted metabolite identification utilized a Thermo model LTQ-Orbitrap Velos mass spectrometer operating in full scan negative ionization MS mode at 60,000 resolution with data-dependent HCD-MS/MS over a mass range of 50-1000 m/z. The mass spectrometer was coupled to a Shimadzu Prominence HPLC system through an electrospray ionization source. Ten microliters of each metabolite extract was injected and separated on a Phenomenex HILIC LC column, 2.0 mm x 100 mm, 3.0 micron particle size, 100 Angstrom pore size, using a gradient of (A) water containing 50 mM ammonium formate, and (B) acetonitrile from 95% to 50% B over 14 minutes as previously descrived. (38). A guard cartridge of matching chemistry was fitted to the analytical column.

### Data analysis

Compound identifications, isotope correction, chromatographic peak alignment, and peak area quantification were performed with MAVEN software (39). Due to the need to simultaneously analyze numerous disparate classes of metabolites in untargeted experiments, only relative quantitation of analytes against a selected internal standard were performed for comparison of values across experimental treatment groups. Multivariate statistical analysis was performed using MetaboAnalyst version 4.0 software (https://www.metaboanalyst.ca/).

### Nutrient downshift survival

*C. jejuni* were grown overnight at 37°C in MH broth under microaerobic condition. Then strains were collected by centrifugation at 6,000 rpm for 10 min and washed twice with MCLMAN minimum media where no carbon and no phosphate present. Strains were diluted to an OD600 of 0.05. After different time intervals, CFU were measured by plating on MH agar plates.

### Osmotic stress survival

*C. jejuni* strains were grown overnight at 37°C in MH broth under microaerobic condition. Then strains were centrifuged and resuspended in MH media containing 1% and 1.5% NaCl. The initial OD of the culture was 0.05 and survival was measured by CFU count at different time intervals (22).

### Real Time PCR

Wild type strain DRH212, *pstS::kan* strains were grown in different condition *in vitro* and total RNA was isolated by the Trizol method and RNeasy Mini Kit (Qiagen). After digestion with TURBO Dnase treatment, cDNA was made by SuperScript III First-Strand Synthesis Supermix (Invitrogen). Then real time PCR was performed by Syber Green method. The gene expression of different targeted genes was measured by 2 ^delta deltaCT^ method.

### Measurement of *in vivo* poly-P levels

Intracellular poly-P levels were measured as described (40) Briefly, strains were grown in MH broth for 24h and 48h. Strains were collected in each time point by centrifugation and dissolved in 250 µL of GITC lysis buffer (4 M guanidine isothiocyanate, 50 mM Tris-HCl, pH 7) and lysed by incubating at 95 °C for 10 min. Lysates were stored at −80 °C. The protein concentration was determined by Bradford assay (Bio-Rad) of a 5-µl aliquot. Poly-P was isolated using EconoSpin silica spin columns (Epoch Life Science) and digested with 1µg of PPX1 exopolyphosphatase from *Saccharomyces cerevisiae*. The resulting free phosphate was measured using a malachite green colorimetric assay and normalized to the total protein (40).

### Chicken colonization assay and *in vivo* phosphate determination

Day-of-hatch white Leghorn chickens were inoculated orally with two doses (approximately 1×10^3^ and 1×10^6^ CFU/ml) of indicated *C. jejuni* strains (wild type strain 81-176, *pstS::kan* and *pstS::kan/C*) (8). Birds were euthanized at day 3 and day 7 after infection, and cecal contents were collected as described (41). Cecal contents were homogenized in sterile PBS and serial dilutions were plated on Mueller-Hinton (MH) agar containing 10% sheep’s blood, cefoperazone (40 µg/ml), cycloheximide (100 µg/ml), trimethoprim (10 µg/ml), and vancomycin (100 µg/ml), selective for *Campylobacter*. Plates were grown for 48 h under microaerobic conditions and colonies were enumerated. For the competition experiment, wild type and *pstS::kan* strain were mixed equally (approximately 1×10^6^ CFU/ml of each strain) and inoculated the day-of-hatch chicks by oral gavage. At Day 7, cecal contents were collected and after serial dilution plated on Campylobacter selective media with and without kanamycin (50 µg/ml). Competitive index values were calculated as previously described (42).

Inorganic free phosphate was measured in chicken cecal content (100mg) at Day 3 and Day 7 by inorganic free phosphate measurement ELISA Kit (Bioassay) as per manufacturer protocol. All chicken experiments and protocols were approved by the Institutional Animal Care and Use Committee in the Animal Care Program at Michigan State University.

## Acknowledgements

We thank Dr. Shannon Manning, Department of Microbiology and Molecular Genetics Michigan State University for providing human stool samples used in this study. This work was supported in part by NIH award AI 111192 (to V.J.D.) and by the Rudolph Hugh Endowment of Michigan State University.

