## Supplementary material for "Phosphate transporter PstSCAB of *Campylobacter jejuni* is essential for lactate-dependent growth and colonization in chickens": Sinha_Pst_Supplemental_Figures (from another version of this preprint: version 1, 2019-11-15)

**Supplementary figures:**

**Fig S1:**

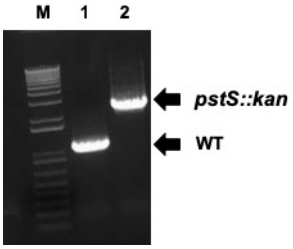


**Fig. S1: Construction of *pstS::kan* insertion mutant:** *pstS::kan* mutant was constructed by insertion of kanamycin cassette into the *pstS* gene of *C.jejuni* strain DRH212. Insertion was confirmed by amplification of *pstS* gene from genomic DNA of DRH212 and kanamycin resistance strains by PCR. Lane M: DNA marker, Lane 1: DRH212; Lane 2: *pstS::kan* mutant.

**Fig S2:**

A)

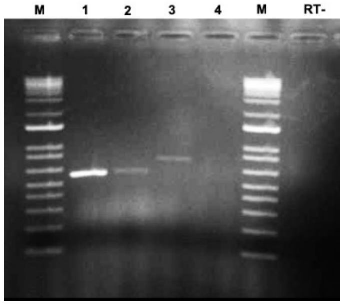


B)

*pstS* Stop codon Intergenic region Start codon

TGGAAAAGTTTTT**TAA**GGAAAAACTTTGTTAAAAGAAAAAATAATCAAATTTGCACTTTTTTT**ATG**TGCT
TTTGTAAGCGTGATAGTAAGTTTTGCTATCATGCTTACAATTTTAATTGAAGCTTTAAAATTCTTTCAAA

*pstC*

***Loop region***

C)

**ΔG=-27.70**

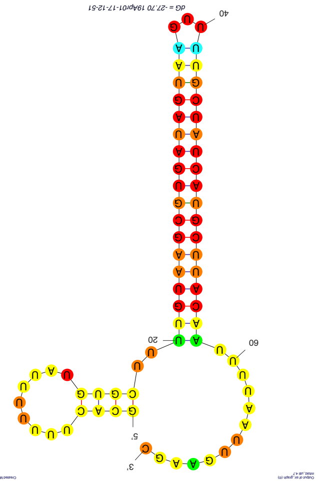

**Fig.S2: A)** Transcript abundance of four genes (*pstS*, *pstC*, *pstA* and *pstB*) of monocystronic *pstSCAB* operon of *C. jejuni* were determined by RT PCR. Wild type strain DRH212 was grown in minimal media containing low phosphate concentration for 6h. Lane 1: *pstS*, Lane 2: *pstC*, Lane 3: *pstA* and Lane 4: *pstB* , RT -: negative control **B)** The underlined region indicated the nucleotide and predicted secondary structure. Stop codon of *pstS* and start codon of *pstC* are indicated in bold. **C)** Secondary structure formed by repetitive palindromic sequence.

**Fig. S3:**

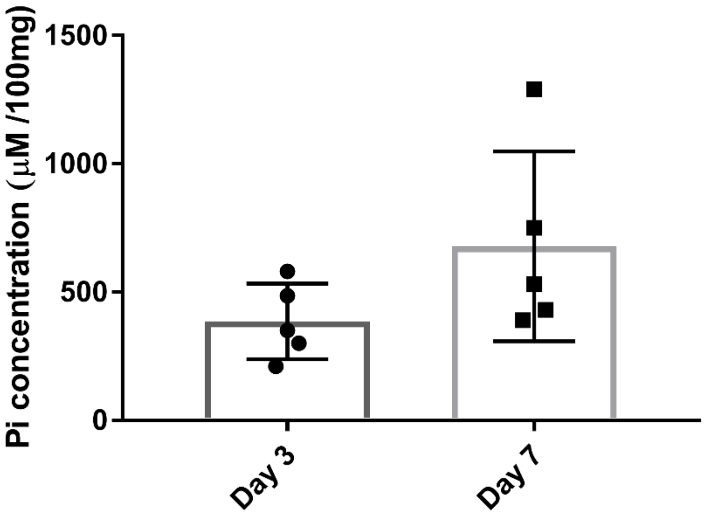


**Fig. S3: Free inorganic phosphate measurement in cecal content of chicken:**

Chicken cecal content were collected from control chicken at day 3 and day 7 and free inorganic phosphate was measured by ELISA.(n=5,±SD).

**Fig. S4:**

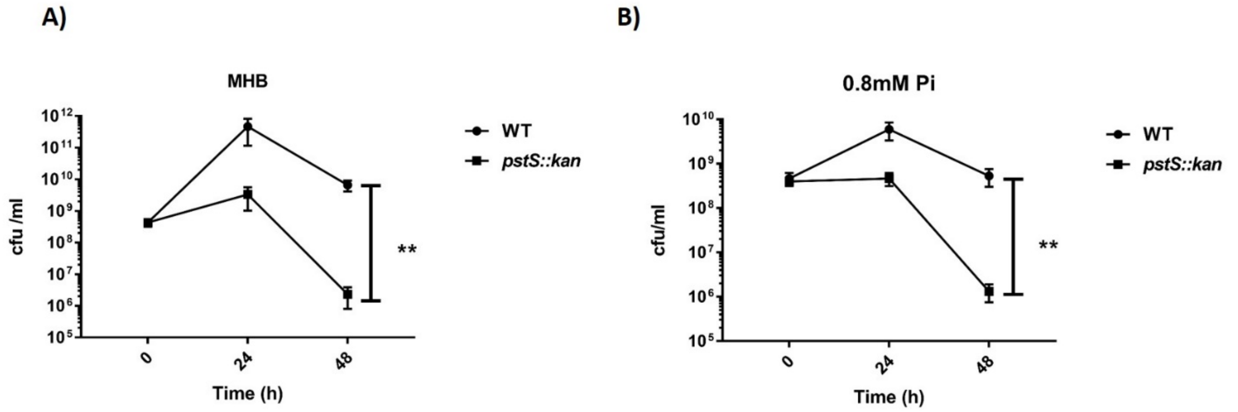


**Fig. S4: *In vitro* competitive growth assay of wild-type and *pstS::kan C. jejuni*.** DRH212 and *pstS::kan* strains were cultured in 1:1 ratio in **A)** nutrient rich (MHB) and **B)** minimal media containing 0.8mM of Pi. Growth of each strain was determined by CFU counts. Statistical analysis was done student t test. $*$ ,P<0.05; $**, P < 0.005$.

**Fig S5:**

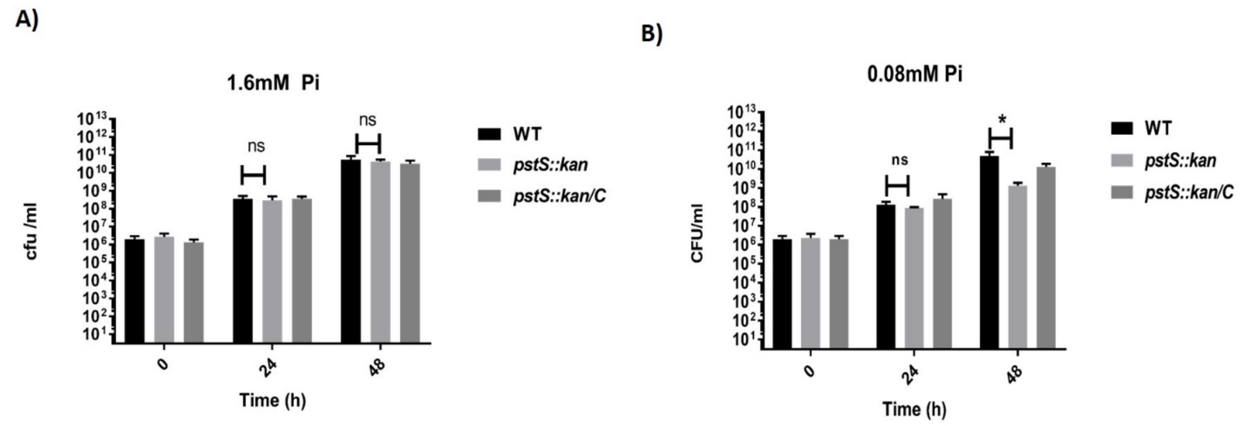


**Fig.S5: Inorganic phosphate is essential for *C. jejuni* growth.** *C. jejuni* strains were grown in minimal media containing pyruvate (10 mM) with two concentrations of inorganic phosphate. Growth of DRH212 and *pstS::kan* was measured by CFU count in minimal media with **A)** 1.6mM Pi and **B)** 0.08mM Pi at 24 and 48h. N=3; error bar indicated SD.Statistical analysis was done by two-way analysis of variation (ANOVAs) with Sidak's multiple comparisions test. $*$ **,P<0.05;** $**, \boldsymbol{P < 0.005}$**.**

**Fig S6:**

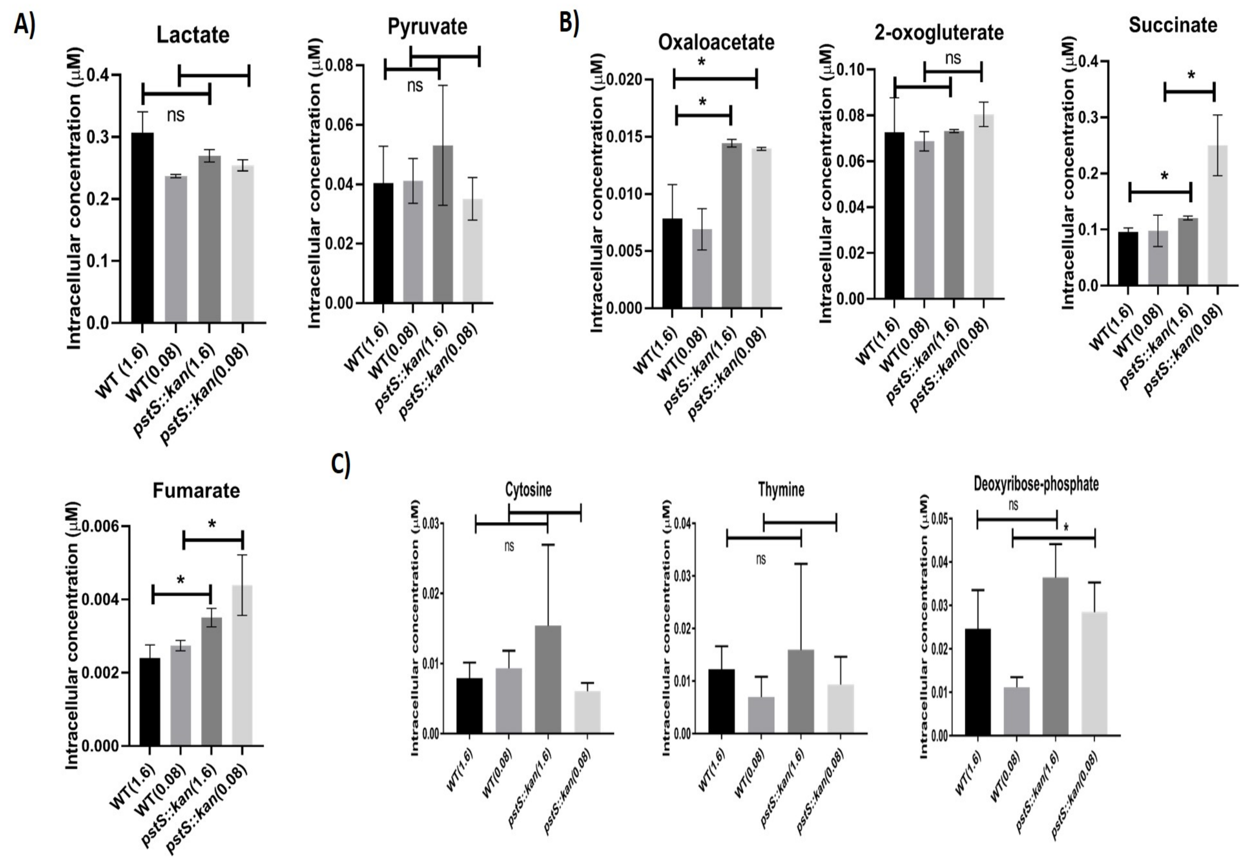


**Fig S6: Metabolites in DRH212 and *pstS::kan* :** DRH212, *pstS::kan* strains were grown in L-lactate minimal media with 1.6 or 0.08 mM Pi at 37°C. After 18h, the intracellular concentration of different metabolites was measured by liquid chromatography mass spectroscopy analysis. **A)** Intracellular concentration of lactate and pyruvate. **B)** Intracellular concentration of TCA cycle intermediates **C)** intracellular concentration of purine, pyrimidine biosynthesis intermediates. All data represents mean value in two independent experiments. The statistical analysis was done by One way ANOVA. $*$ **,P<0.05;** $**, \boldsymbol{P < 0.005}$**.**

**Fig S7:**

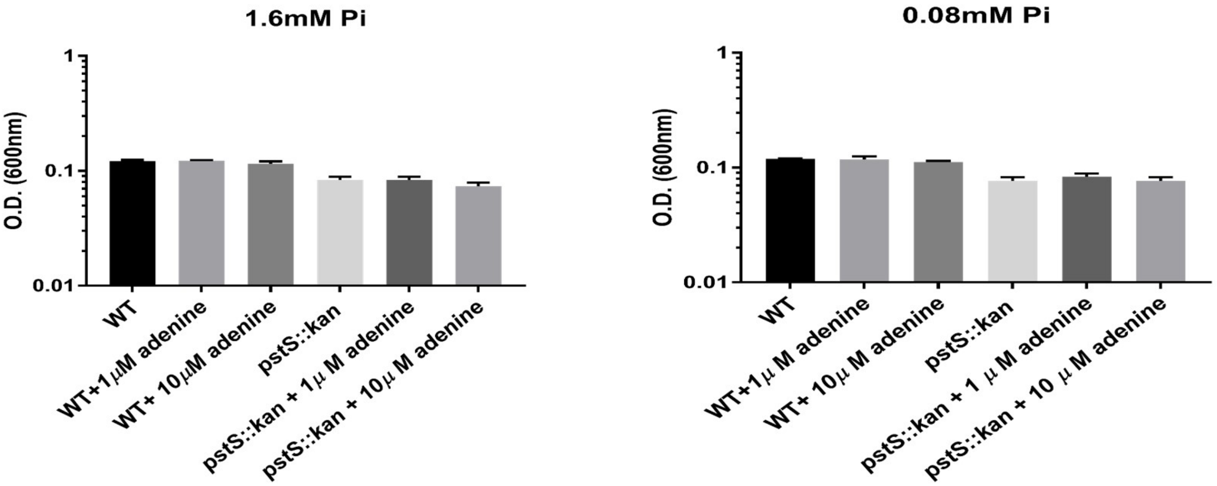


**Fig S7: Adenine supplementation:**

DRH212 and *pstS::kan* grown in different concentration (1μM and 10μM) of adenine were supplemented in L-lactate minimal media with different concentration of inorganic phosphate. After 24h of incubation, O.D at 600nm was measured. N=3; error bar indicated SDs.

**FigS8:**

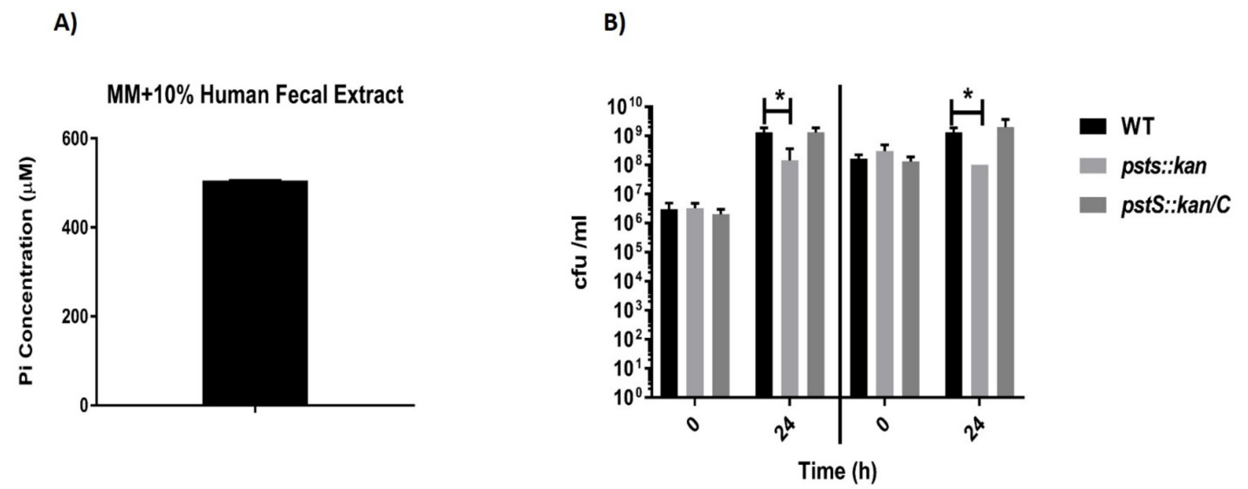


**Fig S8: Inorganic phosphate concentration and growth of *C. jejuni* DRH212 and *pstS::kan* strains in human fecal extract media.** A) Pi concentration of minimal media (no phosphate and no carbon) with 10% of human fecal extracted were measured by ELISA B) Growth of DRH212, *pstS::kan* and *pstS::kan/C* strains were measured in minimal media containing 10% human fecal extract after 24h of incubation by CFUs count. Statistical analysis was done by two-way analysis of variation (ANOVAs) with Sidak's multiple comparisons test. $N = 3$; error bar indicated SDs. $*$, **$P<0.05$; $**$, $P < 0.005$.**
